# Neutrophil-derived complement factor P induces cytotoxicity in basal-like cells via caspase 3/7 activation in pancreatic cancer

**DOI:** 10.1101/2023.10.28.564512

**Authors:** Uday Kishore, Praveen M Varghese, Alessandro Mangogna, Lukas Klein, Mengyu Tu, Laura Urbach, Mengjie Qiu, Remy Nicolle, Valarmathy Murugaiah, Nazar Beirag, Susanne Roth, Dennis Pedersen, Robert B. Sim, Volker Ellenrieder, Gregers Rom Andersen, Roberta Bulla, Shiv K. Singh

**Affiliations:** Department of Veterinary Medicine (CAVM), United Arab Emirates University, Al-Ain, U.A.E; Biosciences, College of Health, Medicine and Life Sciences, Brunel University London, Uxbridge, UK; Institute for Maternal and Child Health, IRCCS (Istituto di Ricovero e Cura a Carattere Scientifico) Burlo Garofolo, Trieste, Italy; Department of Gastroenterology and Gastrointestinal Oncology, University Medical Center, Goettingen, Germany; Department of Surgery, Heidelberg University Hospital, Heidelberg, Germany; Universite Paris, centre de recherche sur l’inflammation /Inserm UMR 1149 Paris, France; Department of Molecular Biology and Genetics, Aarhus University, Gustav Wiedsvej 10C DK8000 Aarhus C, Denmark; MRC Immunochemistry Unit, Department of Biochemistry, University of Oxford, South Parks Road, Oxford OX1 3QU, United Kingdom; Department of Life Sciences, University of Trieste, Trieste, Italy

**Keywords:** Complement, properdin, pancreatic ductal adenocarcinoma, tumour immune microenvironment, neutrophil, apoptosis

## Abstract

Due to profound heterogeneity within stromal immune tumor microenvironment (TME), pancreatic ductal adenocarcinoma (PDAC) remains a hard to treat disease, with the lowest 5-year survival below 10%. Large-scale transcriptomic analysis has revealed two main, clinically relevant PDAC signatures: therapy responsive ‘Classical’ subtype with better prognosis, and poorly-differentiated Basal-like with poor prognosis. It has also become evident that the cellular and humoral components in the immune TME considerably influence the outcome of tumorigenesis. Complement system, a potent humoral innate immune mechanism, also forms a part of this immune TME. In addition to the regular production of various complement components in the liver, certain infiltrating immune cells such as macrophages, dendritic cells and neutrophils, can produce a few complement components locally at the site of infection and inflammation including TME, and modulate tumorigenic outcomes. Neutrophils are the most prevalent innate immune cells in the PDAC TME; however, its role has been attributed as either pro-tumorigenic or anti-tumorigenic. Neutrophils, when stimulated or under stress, are capable of releasing their secretory granules that also contain the only known up-regulator of the complement alternative pathway, Complement Factor P (CFP) or properdin. Properdin can not only facilitate alternative pathway activation by stabilising the C3 convertase, but also act as a pattern recognition receptor on its own and modify inflammatory response. Here, by combining multicenter transcriptome analysis of PDAC patient tumors, single-cell-RNA-seq analysis, preclinical mouse models and human PDAC specimens, we show that properdin expression and neutrophil surveillance are linked to better prognosis in PDAC patients. Furthermore, properdin expression is substantially higher in well-to-moderately differentiated Classical subtype compared to the highly aggressive basal-like PDAC tumours. Mechanistically, exogenous properdin binds to the cell membrane and activates caspase 3/7 to induce apoptosis in basal-like PDAC cells. Together, these findings suggest that the complement protein, properdin, could be a favorable prognostic factor and exhibit anti-tumorigenic functions in PDAC.

## INTRODUCTION

Pancreatic ductal adenocarcinoma (PDAC) remains a challenging disease to treat due to tumour heterogeneity and complex tumour microenvironment (TME) and their molecular cross-talk, with the lowest 5-year survival below 10% (Sung et al., 2021; Ushio et al., 2021). In comparison to other cancer types, immunotherapy has does not appear to be very effective in PDAC patients (Pihlak et al., 2018; Riquelme et al., 2018; Huber et al., 2020; Timmer et al., 2021). PDAC TME exhibits complex stromal immune heterogeneity, which is one of the major reasons for poor therapeutic outcome in patients (Huber et al., 2020).

Genetically, PDAC patients harbour alteration or amplification of *KRAS*, followed by deletion or inactivating mutation of *SMAD4* and *TP53* (Jiang et al., 2022b). KRAS is a member of the RAS family of genes that code for small GTPases, which are crucial for cell proliferation, differentiation, and survival (Downward, 2003). Approximately 90- 95% of PDAC tumours harbour oncogenic single amino acid substitution mutations in KRAS at either G12, G13, or Q61 (Almoguera et al., 1988; Rozenblum et al., 1997; Löhr et al., 2005; Jones et al., 2008). These mutations result in constitutive activation of the KRAS protein, leading to continuous signal transduction and uncontrolled cellular proliferation (Scheidig et al., 1999). Given its high prevalence, oncogenic KRAS mutation is considered an early and critical event in the development of PDAC (Bailey et al., 2016). SMAD4, is a central mediator of the transforming growth factor-beta (TGF-β) signaling pathway, which is involved in cellular processes such as growth suppression, differentiation, and apoptosis (Massague et al., 2000; Bardeesy et al., 2006). Inactivation of SMAD4, either through mutations or deletions, is observed in about 90% of PDAC cases (Hahn et al., 1995; Hahn et al., 1996; Grant et al., 2016). Loss of functional SMAD4 leads to a disruption of TGF-β signaling, which contributes to tumor progression, metastasis, epithelial-to-mesenchymal transition (EMT) and resistance to certain therapies (David et al., 2016; Grant et al., 2016). TP53 is a tumor suppressor gene that encodes the p53 protein (Grant et al., 2016). The p53 protein plays a pivotal role in responding to cellular stress, such as DNA damage, by inducing cell cycle arrest, apoptosis, or senescence to prevent the propagation of genetically damaged cells (Wang et al., 2021). Mutations in TP53 are observed in approximately 50-70% of PDACs (Christenson et al., 2020; Mizrahi et al., 2020). The inactivation or dysfunction of p53 in PDAC allows cells with DNA damage to evade apoptosis and continue to proliferate, leading to genomic instability and tumor progression (Christenson et al., 2020; Mizrahi et al., 2020; Wang et al., 2021). Besides genetic alterations, large-scale transcriptome profiling identified two main PDAC subtypes in PDAC patients: the classical (CLA) and basal-like (BL) (Moffitt et al., 2015; Bailey et al., 2016; Miyabayashi et al., 2021). The CLA subtype is associated with better prognosis, whereas the BL subtype is linked to worse clinical outcome in PDAC patients (Moffitt et al., 2015; Nicolle et al., 2017). However, how TME-derived factors or other immune cell types (e.g. neutrophils and macrophages) influence PDAC subtype plasticity and patients prognosis is not fully understood.

A crucial contribution to the innate immune surveillance against cancer comes from the complement system. A bioinformatic analysis of 30 different cancer types revealed that breast cancer, lung adenocarcinoma, liver hepatocellular carcinoma, cervical squamous cell carcinoma, PDAC, prostate adenocarcinoma, mesothelioma, sarcoma, skin cutaneous melanoma have favourable prognosis when they are associated with high expression of complement genes (Mangogna et al., 2019; Mangogna et al., 2020). However, PDAC cells have also been demonstrated to exhibit resistance to complement attack due to higher levels of inhibitory anti-proteases alpha-2-macroglobulin and C1INH in the serum; overexpression of TGF-β is also known to inhibit C3 secretion in PDAC cells (Bettac et al., 2017).

The complement activation can take place via three pathways: classical, alternative, and lectin, depending upon the target ligands and their recognition subcomponents (Lu and Kishore, 2017). Properdin, a ∼50 kDa glycoprotein found at 4–25 μg/ml level in plasma, is the only known positive regulator of the alternative pathway (Pangburn, 1989; Chen et al., 2018). Properdin is also known to act as a humoral pattern recognition receptor (PRR) for ligands such as lipopolysaccharide, acetylated low-density lipoprotein, zymosan and apoptotic cells (Chen et al., 2018; El-Ghobarey and Whaley, 1980; Kouser et al, 2013). Recent studies showing properdin function as a pattern recognition receptor (PRR) and modulator of pro-inflammatory immune response have suggested its importance role beyond the upregulation of the complement alternative pathway (Al-Mozaini et al., 2018; Kouser et al., 2018). Additionally, properdin is also locally produced in secondary granules of stimulated peripheral blood neutrophils, DCs and T cells (Schwaeble et al., 1993; Wirthmueller et al., 1997; Vuagnat et al., 2000; O‘Flynn et al., 2013; Dixon et al., 2017). Since up to 80-90% of the terminal pathway is activated via the alternative pathway and properdin is known to modulate the pro-inflammatory response, the role of properdin in malignancy has been examined. Analysis of gene mutations between primary breast tumours and normal tissue revealed mutations in the properdin (*CFP*) gene (Sjöblom et al., 2006). In breast cancer, properdin has been shown to inhibit tumor proliferation, independent of complement-mediated mechanisms, by upregulating the intracellular pro-apoptotic transcription factor, DNA damage-inducible transcript 3 (DDIT3), which is intrinsically associated with the endoplasmic reticulum-stress response (Block et al., 2019).

In the current study, we analysed the protein and mRNA levels of properdin in PDAC and interrogated if properdin could be a surrogate marker of disease severity and overall survival in PDAC. Moreover, we investigated the correlation of properdin with tumour-infiltrating immune cells (TIICs), especially neutrophils, in PDAC. We also performed immunohistochemical analysis to identify the most likely source(s) of properdin in the PDAC TME. Subsequently, we examined the effects of exogenous properdin treatment on PDAC cells.

## MATERIALS AND METHODS

### Single-cell RNA sequencing analysis

Raw single-cell RNA sequencing data was downloaded from the Gene Expression Omnibus (GEO) database with the accession ID GSE205049, GSE155698 and GSE212966. Data were then pre-processed and clustered separately, following the pipeline as described previously (Yousuf et al, 2023; Steele et al, 2020; Chen et al, 2023). Cells with more than 25% mitochondrial genes and fewer than 100 genes were removed. Raw counts were normalized and then scaled to regress out the percentage of mitochondrial genes and the counts of genes. Principal component analysis and the SNN graph construction were implemented with the embedded functions in Seurat 4.0. Canonical markers and gene profiles were referred to annotate cell populations in each dataset. The normalized complement factor P/properdin (*CFP)* expression level was then used to generate heatmaps.

### Properdin gene expression analysis in PDAC

The expression of the *CFP* gene between PDAC and their normal tissue counterparts was analyzed using the GEPIA2 (Gene Expression Profiling Interactive Analysis 2). GEPIA2 is a comprehensive interactive web resource for analyzing cancer OMICS data based on TCGA (The Cancer Genome Atlas) and GTEx (Genotype-Tissue Expression) data (Tang et al., 2019). GEPIA2 is available via http://gepia2.cancer-pku.cn/. GEPIA2 generates box plots in order to compare the differences in mRNA levels between normal tissue and PDAC. To analyse the mRNA expression levels of *CFP* in the neoplastic and healthy tissues, *p*-value ≤0.01 and induction or repression ratios higher than Log2-fold change (Log2FC) >1 criterion were applied to identify statistically significant differential gene expression. The differential analysis was based on the selected datasets [TCGA-PDAC *vs* normal tissue (TCGA pancreas + GTEx pancreas)]. TCGA-PDAC contains 179 pathological samples whereas TCGA pancreas + GTEx pancreas had 171 normal tissues. The method for differential analysis was one-way ANOVA. The Log2[transcripts per million (TPM) + 1] transformed expression data was used for plotting.

### Properdin protein expression analysis in PDAC

The differential protein expression of properdin between 137 PDACs and 74 normal pancreatic tissues was analyzed via UALCAN, a comprehensive interactive web resource for analyzing cancer OMICS data, which allowed protein expression analysis using the Clinical Proteomic Tumor Analysis Consortium (CPTAC) Confirmatory/Discovery dataset (Chen et al., 2019; Zhang et al., 2022). The web server is publicly accessible via http://ualcan.path.uab.edu/analysis-prot.html. The CPTAC has extensive mass spectrometry-based proteomics data for PDAC from the respective TCGA. Z-value represents standard deviation from the median across samples for PDAC. Log2 Spectral count ratio values from CPTAC were first normalized within each sample profile, then normalized across samples. *p*-value ≤0.05 was considered statistically significant.

### Tumour-infiltrating neutrophil in PDAC

Levels of tumor-infiltrating neutrophils in PDAC were analyzed from TCGA-PDAC dataset using Tumor Immune Estimation Resource (TIMER) version 2.0. TIMER2 is a comprehensive resource for systematic analysis of immune infiltrates across diverse cancer types (Li et al., 2017). TIMER2 is available via http://timer.cistrome.org/. The gene module allows visualization of the correlation between *CFP* expression and neutrophil infiltration level in PDAC. The scatter plot generated presents the relationship between infiltrate estimation value and gene expression. Tumor purity is a major confounding factor in this analysis since most immune cell types are negatively correlated with tumor purity. Hence, the “Purity Adjustment” option was used, which used the partial Spearman’s correlation to perform association analysis. In TIMER2, the correlation of gene expression was evaluated by Spearman’s correlation and statistical significance (*p*-value ≤0.05 was considered statistically significant). The strength of the correlation was determined using the following guide for the absolute value: 0.00-0.19: very weak; 0.20-0.39: weak; 0.40-0.59: moderate; 0.60-0.79: strong; and 0.80-1.0: very strong.

### *In silico* analysis of survival in PDAC

The prognostic significance of *CFP* mRNA expression with respect to survival in PDAC was analyzed. Survival curves were generated by TIMER2 and GEPIA2 using genomic data from TCGA-PDAC to generate survival probability plots via Kaplan– Meier (KM) survival analysis (Li et al., 2017; Tang et al., 2019). Hazard ratio (HR) with a 95% confidence interval and Logrank *p*-value were also computed. *p*-value ≤0.05 or Logrank *p*-value ≤0.05 were considered statistically significant. The HR and *p*-value for Cox proportional hazard model and the Logrank *p*-value for KM curve are shown on the KM curve plot.

### Patient data heatmaps

Normalised gene expression values of the Puleo (Puleo et al., 2018), TCGA (Cancer Genome Atlas Research Network. Electronic address and Cancer Genome Atlas Research, 2017), Moffitt (Moffitt et al., 2015), Bailey (Bailey et al., 2016) and PDX (Nicolle et al., 2017) cohorts were classified using the Microenvironment Cell Populations counter tool, MCPcounter, to determine neutrophil scores of patients (Becht et al., 2016).

For CLA-related gene signatures, the average expression of the gene-wise centred (non-scaled) expression values were used to calculate Gene Set Correlation Enrichment (Gscore; http://140.113.120.31/Gscore_test_hua/index.php) for the TCGA (Cancer Genome Atlas Research Network. Electronic address and Cancer Genome Atlas Research, 2017), Moffitt (Moffitt et al., 2015), Bailey (Bailey et al., 2016), PDX (Nicolle et al., 2017) and Notta (Chan-Seng-Yue et al., 2020) signatures. For the Puleo signature, the projection of the original publication was used (Puleo et al., 2018).

Heatmaps were generated in R v4.1.2 using the heatmap package v1.0.12. Values were normalised using the scale = “row” parameter and clustered by column. The hierarchical clustering dendrogram was separated into potentially biologically relevant clusters using the cutree function. For “Neutrophil Group” and “PDAC Signature Group” annotation, patients were ranked by the respective MCPcounter or Gscores and separated into the lower (Q1), middle (Q2) and upper quartile (Q3).

For CFP expression, RMA-normalized probe expression values of the original publication were used (Puleo et al., 2018). For survival analysis, heatmap clustering information was extracted using cutree. The corresponding patient overall and disease-free survival information (as given in the original publication) (Puleo et al., 2018) was plotted as Kaplan-Meier survival curve per cluster.

### Mouse and human PDAC tissues

The *Kras^G12D^; p53^R172H^; Cre* (KPC) mouse model is an established and clinically relevant model of PDAC. The PDAC development and progression in KPC mice closely mirrors that of human pathology. The model replicates a number of the key histological (such as cellular morphology, poor vascularity, fibrosis, and metastatic spread) and clinical (such as ascites development, bowel and biliary obstruction, and cachexia) characteristics associated with human PDAC (Hingorani et al., 2005; Olive et al., 2009; Beatty et al., 2011; Provenzano et al., 2012; Feig et al., 2013; Jacobetz et al., 2013; Lee et al., 2016). A substantial inflammatory response and the exclusion of effector T cells are two other important aspects of the immune TME that the model also replicates (Beatty et al., 2011; Bayne et al., 2012; Keenan et al., 2014; Beatty et al., 2015; Lee et al., 2016). As a result, the KPC model is a useful tool for dissecting the biological mechanisms of inflammatory cell recruitment and T cell exclusion (Lee et al., 2016). Tissues samples were collected from KPC mice under the approval of the University Medical Center Göttingen (UMG) Central Animal-experimental Facility. The in-house pathologist conducted histopathological grading of KPC tumour tissues. Human primary PDAC tissues were obtained from the Institute of Pathology, UMG.

### Immunofluorescence staining

Immunofluorescence staining was performed, as described previously (Tu et al., 2021). Briefly, tissue sections were deparaffinised in xylene twice for 30 mins, each time prior to rehydration in ethanol. Next, sections were kept in citrate buffer (pH 6.0) for 6 mins in the microwave for antigen retrieval. Sections were then washed five times with PB buffer and blocked in 10 % normal goat serum (NGS; Abcam) at 4°C for 1.5 h. Primary monoclonal antibodies to human properdin (Santa Cruz; 1:100) and amylase (Cell Signaling Technology; 1:100) were diluted with 2 % of NGS, added onto the sections, and incubated at 4°C overnight. After washing with PB buffer five times, sections were incubated with donkey anti-rabbit-Alexa Fluor® 488 (ThermoFisher Scientific, 1:500) and donkey anti-mouse-Alexa Fluor® 568 (ThermoFisher Scientific, 1:500) conjugates at 4°C for 2h, followed by DAPI staining. Images were obtained via FluoView 1000 confocal microscope (Olympus).

### Properdin Purification

Recombinant human properdin was expressed and purified, as previously described (Pedersen et al., 2017). Briefly, human properdin cDNA was subcloned into the pCEP4 mammalian expression vector. HEK293F cells in suspension used for expression were maintained in serum-free FreeStyle 293 Expression Medium (Invitrogen) at 37°C, 8% CO2, 125 rpm. Polyethyleneimine (PEI) and 1 mg/l plasmid DNA were used to transiently transfect cells. 3–4 days after transfection, the conditioned medium was collected, diluted two-fold with water, and adjusted to pH 6.4. A human C3b-Sepharose column equilibrated with 20 mM imidazole, 50 mM NaCl, 5 mM EDTA, pH 6.4 was used to purify secreted properdin. The bound protein was eluted with 15 bed volumes of imidazole buffer containing 500 mM NaCl and concentrated to about 0.5 mg/ml. Properdin was dialyzed against 50 mM phosphate buffer, 2.5 mM EDTA, pH 6.0, prior to cation exchange chromatography. The protein solution was loaded on to a 1 ml Mono S column (GE Healthcare), eluted with a 20-bed volume linear gradient from 0 to 500 mM NaCl, and dialyzed against 10 mM HEPES, 75 mM NaCl, pH 7.2 buffer.

### Properdin binding and apoptosis studies on PDAC cell lines

PANC-1 (CRL-1469) and MiaPaCa-2 (CRL-1420) cells were obtained from ATCC and subjected to Immunofluorescence staining. PANC-1 and MiaPaCa-2 were cultured and maintained in growth medium [DMEMF/12 with GlutaMAX™ Supplement (Gibco), + 10% v/v Heat Inactivated Fetal Bovine Serum (FBS) (Gibco) + 1% v/v Penicillin/Streptomycin (PS) (Gibco)] at 37°C under 5% v/v CO2. For the binding experiments, PANC1 and MiaPaCa-2 cells (0.4 × 10^6^ cells/well) were seeded on collagen-coated coverslips in a 6 well plate. Both cell lines were serum-starved for 24h using serum-free DMEMF/12 with GlutaMAX™ Supplement + 1% v/v PS. For properdin binding analysis, the cells were treated with human properdin (20 µg/ml) for 1h at 37 °C, followed by 1 h incubation at 4°C. The coverslips were incubated for 1 h with rabbit anti-human properdin polyclonal antibodies (MRC Immunochemistry Unit, Oxford; 1:200). Following washes with PBS, the cells were incubated with goat anti-rabbit IgG conjugated to Alexa Fluor® 488 (1:200) (Abcam) and Hoechst (1:10,000) (Sigma-Aldrich). Similarly, for apoptosis analysis, serum-starved PANC-1 or MiaPaCa-2 cells were incubated with properdin (20 µg/ml) for 24 h and 48 h, respectively. Post-treatment, the cells were incubated with annexin V binding buffer containing FITC annexin V (1:200) (Biolegend), propidium iodide (PI) (1:200) (Biolegend), and Hoechst (1:10,000) for 1h at room temperature (RT) in the dark. After washing with PBS three times, the coverslips were mounted on slides and viewed under an HF14 Leica DM4000 microscope.

### MTT assay

PANC-1 and MiaPaCa-2 (0.1 × 10^5^) cells were seeded in a 96-well microtiter plate. Serum starved PANC-1 and MiaPaCa-2 cells were incubated with human properdin (20 µg/ml) for 24 h and 48 h, respectively. Next, MTT (3-[4,5-dimethylthiazol-2-yl]-2,5-diphenyltetrazolium bromide) (Thermo Fisher) assay was performed by incubating the cells with 50 µg/µl MTT (5 mg/ml stock) per well for 4 h at 37°C. All but 25 µl of media per well was removed, and 50 µl of dimethyl sulfoxide was added to each well. The plate was incubated for another 10 min at 37°C, and the absorbance was read at 570 nm using a Bio-Rad plate reader.

### Flow Cytometry

PANC-1 and MiaPaCa-2 (0.4 × 10^6^) cells were seeded in a six-well-plate. Serum starved PANC-1 and MiaPaCa-2 cells were incubated with human properdin (20 µg/ml) for 24 h and 48 h, respectively. The cells were trypsinised and incubated with annexin V binding buffer containing FITC annexin V (1:200) and PI (1:200) for 1h at RT in the dark. After washing with PBS twice, apoptosis was measured using a NovoCyte flow cytometer. Compensation parameters were acquired using unstained, FITC-stained only, and PI-stained only cells.

### Caspase3/7 activation assay

PANC-1 and MiaPaCa-2 cells (0.1 × 10^5^) were grown in a 96-well microtiter plate. Serum starved PANC-1 and MiaPaCa-2 cells were incubated with human properdin (20 µg/ml) and CellEvent™ Caspase-3/7 Green Detection Reagent (Invitrogen) (5 μM) for 48 h. Staurosporine (1 μM; Sigma-Aldrich) was used as a positive control for apoptosis. The kinetic assay was performed by incubating the plate at 37 °C in a CLARIOstar Plus plate reader (BMG Labtech) for the duration of the study. Fluorescence corresponding to Caspase-3/7 activation was recorded every hour using the plate reader’s FITC/Alexa Fluor™ 488 filter settings.

### Caspase 8/9 activation assay

PANC-1 and MiaPaCa-2 cells (0.1 × 10^5^) were grown in a 96-well microtiter plate. Serum starved PANC-1 cells were incubated with human properdin (20μg/ml)) for various time points (3h, 6h, 12h). Staurosporine (1 μM) was used as a positive control for apoptosis. Similarly, serum-starved MiaPaCa-2 cells were incubated with properdin (20μg/ml)) for 6h, 12h and 18h. Post-treatment, 100 µl of either Caspase-Glo® 8, or 9 Reagent (Promega) with MG-132 Inhibitor (Promega) was added to each well. The plate was incubated for another 30 mins at RT. Luminance corresponding to Caspase-8/9 activation was recorded using a CLARIOstar Plus plate reader (BMG Labtech).

## RESULTS

### Infiltrating neutrophils produce properdin in PDAC tumour microenvironment

To determine the source of *CFP* expression in the pancreatic exocrine, endocrine, neoplastic, stromal as well as immune cells, we comprehensively analysed scRNA-seq both in adjacent normal (AdjNorm) and pancreatic tumour tissues of PDAC patients (**Fig. 1A**). The *CFP* expression was mainly enriched in the neutrophil, dendritic cells and monocytes among the three distinct datasets. We also noted weaker expression of CFP in the AdjNorm tissues. Although the number of samples was relatively small, it is evident that the immune cells are the major cell type expressing CFP, rather than neoplastic cells (**Fig. 1A**). We next sought to examine whether CFP expression determined patient prognosis and overall clinical outcome of PDAC patients. Hence, we analysed mRNA and protein expression of CFP (properdin) in normal and tumour tissues by using UALCAN tool in a large cohort of PDAC datasets such as TCGA (**Fig. 1B and C**). We noted a significant increase in properdin mRNA expression in PDAC tumors compared to the normal pancreas (**Fig. 1B**). Similarly, UALCAN proteomics analysis revealed a higher expression of properdin in PDAC than the corresponding healthy pancreas tissue (**Fig. 1C**).

**Figure 1.**
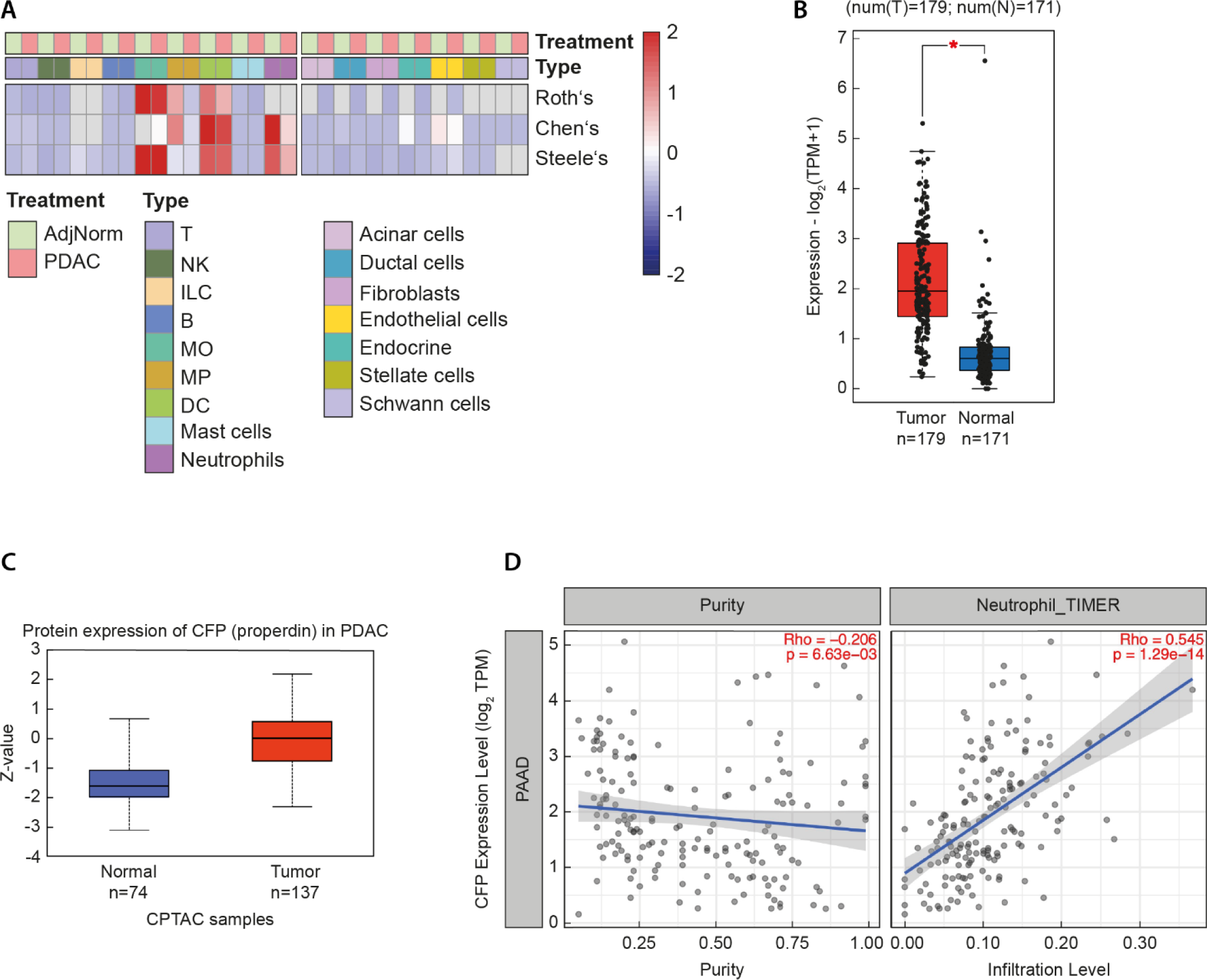
Infiltrating Neutrophils release properdin in PDAC microenvironment. (A) Heatmap of the *CFP* expression levels in each cell population from three published datasets indicate a consistent expression pattern (Yousuf et al, 2023; Steele et al, 2020; Chen et al, 2023). Grey boxes indicate undetected cell populations in the corresponding dataset. **(B)** A higher expression of *CFP* was detectable in PDAC (red) compared to normal pancreas tissue (blue). (* *p*-value <0.01). TPM, transcripts per million. **(C)** Higher levels of properdin in PDAC compared to healthy tissues, similar to the gene expression studies. (*p*-value <0.0001). [Normal (n=74 samples); Maximum: 0.667, Upper quartile: -1.105, Median: -1.618, Lower quartile: -2.006, Minimum: - 3.124; Tumor (n=137 samples); Maximum: 2.191, Upper quartile: 0.553, Median: 0.009, Lower quartile: -0.76, Minimum: -2. 323]. **(D)** *CFP* expression shows a significant negative correlation with tumor purity, whereas significantly positively correlating with levels of infiltrating neutrophils. [r = 0.545; p = 1.29 x 10-14]. Genes highly expressed in the tumor microenvironment are expected to have negative association with tumor purity, while the opposite is likely to be the case for genes highly expressed in the tumor cells.

In order to ascertain if properdin found in PDAC tissue was secreted/produced by PDAC cells or another component of the TME, the tumour purity was analysed. Tumour purity refers to the proportion of cancer cells in the TME. Genes highly expressed in the TME are expected to have a negative association with tumor purity, while the converse is likely to be case for the genes highly expressed in the tumor cells. Based on the TIMER V2.0 analysis, CFP expression showed a significantly negative correlation with tumour purity (**Fig. 1D**) [r = -0.206; p = 6.63 x 10^-3^]. This negative correlation observed between CFP expression levels and tumour purity indicated that PDAC cells did not produce CFP. Neutrophils are among the most abundant immune cell type within the PDAC TME; when stimulated or under stress, they are known to release properdin stored in their secondary granules. Thus, a correlation between CFP gene expression and neutrophil infiltration level in PDAC was assessed via TIMER V2.0 analysis. It revealed a significant and moderate positive correlation with infiltrating levels of neutrophils (**Fig. 1D**). Thus, the results of the TIMER V2.0 analysis suggested that infiltrated neutrophils were a major source of properdin found in the PDAC TME.

### High properdin expression level associates with improved survival of PDAC patients

Kaplan–Meier survival analysis using TCGA-PDAC dataset (n=177) was performed to establish an association between CFP expression and overall survival (OS). The OS of PDAC patients expressing high and low levels of properdin over a period of 50 months was analyzed by TIMER2. The analysis revealed that patients with high levels of properdin expression in the TME showed a significant improvement in survival compared to their low expression counterparts (**Fig. 2A**) Similarly, OS of BL or CLA subtype patients expressing high and low levels of properdin over a period of 40 or 80 months, respectively, was analyzed with Kaplan–Meier survival analysis using TCGA-PDAC dataset [BL subtype patients (n=32); CLA subtype patients (n=42)] by GEPIA2. In case of both BL subtype patients (**Fig. 2B**), or CLA subtype patients (**Fig. 2C**), OS was found to be significantly improved in patients with high levels of properdin expression compared to their low expression counterparts. Finally, the cumulative survival analysis for CFP expression and levels of infiltrating neutrophils in PDAC over a period of 60 months was also assessed by analyzing the TCGA-PDAC dataset (n=178) by Kaplan-Meier survival analysis using TIMER2 (**Fig. 2D**). The analysis revealed that patients with higher levels of infiltrating neutrophils but reduced CFP expression were associated with a poorer OS. PDAC microarray data (Puleo et al., 2018) revealed that most patients with a high properdin expression also had a higher infiltration of neutrophils, as determined by MCPcounter (**Fig. 2E**; cluster 1). Crucially, this particular cluster of patients with high CFP expression and high neutrophil infiltration had a trend towards an improved overall and disease-free survival (**Fig. 2F and G**). However, the improved overall and disease-free survival was not observed for patients with either high neutrophil scores but low CFP expression (cluster 2), or patients with both low neutrophil score and low CFP (cluster 3; **Fig. 2F**). These results reaffirmed that properdin mRNA expression positively correlated with the OS rate in PDAC patients.

**Figure 2.**
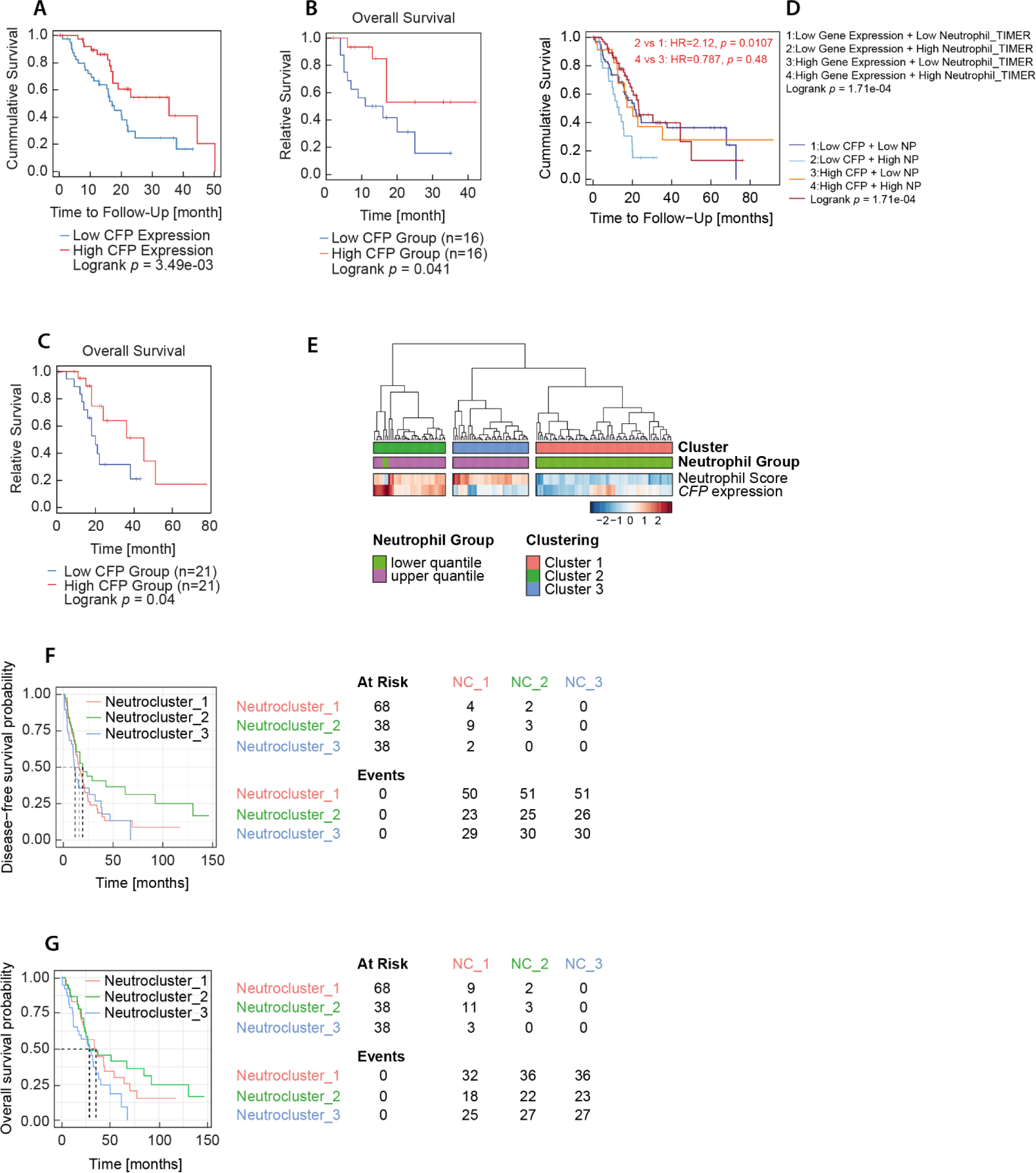
Properdin expression associates with PDAC patient prognosis. (A) Overall survival (OS) of PDAC patients expressing high [red] (n=45) and low [blue] (n=132) levels of properdin over a period of 50 months analyzed with Kaplan–Meier survival analysis using the publicly available TCGA-PDAC dataset (n=177). Patients with high levels of properdin expression show significant improvement in survival compared to their low expression counterparts. [Kaplan-Meier Curve Parameters: Split Expression Percentage of Patients: 25%; Group Cutoff: quartile; Cutoff-High = 75% (high CFP expression: n = 45); Cutoff-Low = 25% (low CFP expression: n = 132)]. **(B)** OS of BL subtype patients expressing high [red] (n=16) and low [blue] (n=16) levels of properdin over a period of 40 months analyzed with Kaplan–Meier survival analysis using TCGA-PDAC dataset (n=32). Patients with high levels of properdin expression have a significantly improved survival compared to their low expression counterparts. [Kaplan-Meier Curve Parameters: Split Expression Percentage of Patients: 25%; Group Cutoff: quartile; Cutoff-High = 75% (high CFP expression: n = 16); Cutoff-Low = 25% (low CFP expression: n = 16)]. **(C)** OS of CLA subtype patients expressing high [red] (n=21) and low [blue] (n=21) levels of properdin over a period of 80 months analyzed with Kaplan–Meier survival analysis using TCGA-PDAC dataset (n=42). Patients with high levels of properdin expression have a significant improvement in survival compared to their low expression counterparts. [Kaplan-Meier Curve Parameters: Split Expression Percentage of Patients: 25%; Group Cutoff: quartile; Cutoff-High = 75% (high CFP expression: n = 21); Cutoff-Low = 25% (low CFP expression: n = 21)]. **(D)** OS of PDAC patients was evaluated by combining CFP expression and levels of infiltrating neutrophils (TIMER2) over a period of 60 months analyzed by Kaplan-Meier survival analysis using TCGA-PDAC dataset (n=178). All the conditions have statistical significance in the correlation with patients’ survival (Logrank test *p* =1.71e-04): low CFP expression/low neutrophils (blue); low CFP expression/high neutrophils (light blue); high CFP expression/low neutrophils (orange); high CFP expression/high neutrophils (red). Assuming low CFP expression as an independent variable, higher levels of infiltrating neutrophils correlated with a poor prognosis (HR=2.12; *p*=0.0107), whereas no statistically significant difference was observed for high CFP expression while considering low or high neutrophils’ levels. [Kaplan-Meier Curve Parameters: Split Expression Percentage of Patients: 50%; Split Infiltration Percentage of Patients: 50%]. **(E)** Patients of the Puleo cohort with MCPcounter neutrophil scores in the upper and lower quartile in this cohort (n=77 each) were plotted for neutrophil scores and CFP expression. Row-normalised values are shown. (**F) and (G**) Clustering information of E was extracted, and per- and disease-free survival (DFS) **(F)** cluster overall survival (OS) (OS) **(G)** were plotted. Cluster 1, n=39; cluster 2, n=41; cluster 3, n=74.

### Neutrophil infiltration is associated with CLA PDAC differentiation

As mentioned previously, transcriptomic subtyping has become increasingly important in PDAC. Hence, we investigated whether the favourable neutrophil-properdin axis is correlated with the phenotypic tumour subtype identity. Utilising classifications from the Puleo, Notta, Moffitt and Bailey cohorts and PDX analysis, we interrogated the relationship between the CLA phenotype and neutrophil infiltration (as determined by MCPcounter). As properdin expression and neutrophil infiltration were found to confer improved OS, we hypothesised an association of the neutrophil score with the CLA signatures. Indeed, patients with low infiltration of neutrophils generally showed low enrichment of CLA signatures (**Fig. 3A**). Although patients with low neutrophil infiltration may also show CLA differentiation, a high CLA score was associated with a high neutrophil score. Across all these signatures, patients in the higher neutrophil score quartile had significantly higher CLA signature scores (**Suppl. Fig. 1**). Next, we sought to examine properdin expression in CLA and BL tumours. In our earlier studies, we have shown that *Kras^G12D^; p53^R172H^; Cre* (KPC)-derived tumours clearly exhibited low-grade/CLA and high-grade/BL tumours with GATA6^low^ and GATA6^high^, respectively (Tu et al., 2021; Krebs et al., 2022) The presence of properdin in the PDAC TME was evaluated using immunofluorescence staining of properdin and amylase in those KPC-derived tumours, as previously described (Hingorani et al., 2005; Tu et al., 2021). In line with the results of Fig. 3A, we found strong expression of properdin in the low-grade (G1 and G2)/CLA tumours compared to high-grade (G3 and G4)/BL tumors derived from KPC mice (**Fig. 3B and C**). Furthermore, we also noted high expression of properdin in low-grade/CLA-like compared to high-grade/BL human PDAC tumors (**Fig. 3D and E**). Together with patient survival data, immuno-staining of murine and human PDAC tissues confirmed that properdin was mainly produced in the low grade or W/M tumours compared to high grade or poorly differentiated PDAC tissues, suggesting a negative correlation between neutrophils and the aggressive PDAC phenotype.

**Figure 3.**
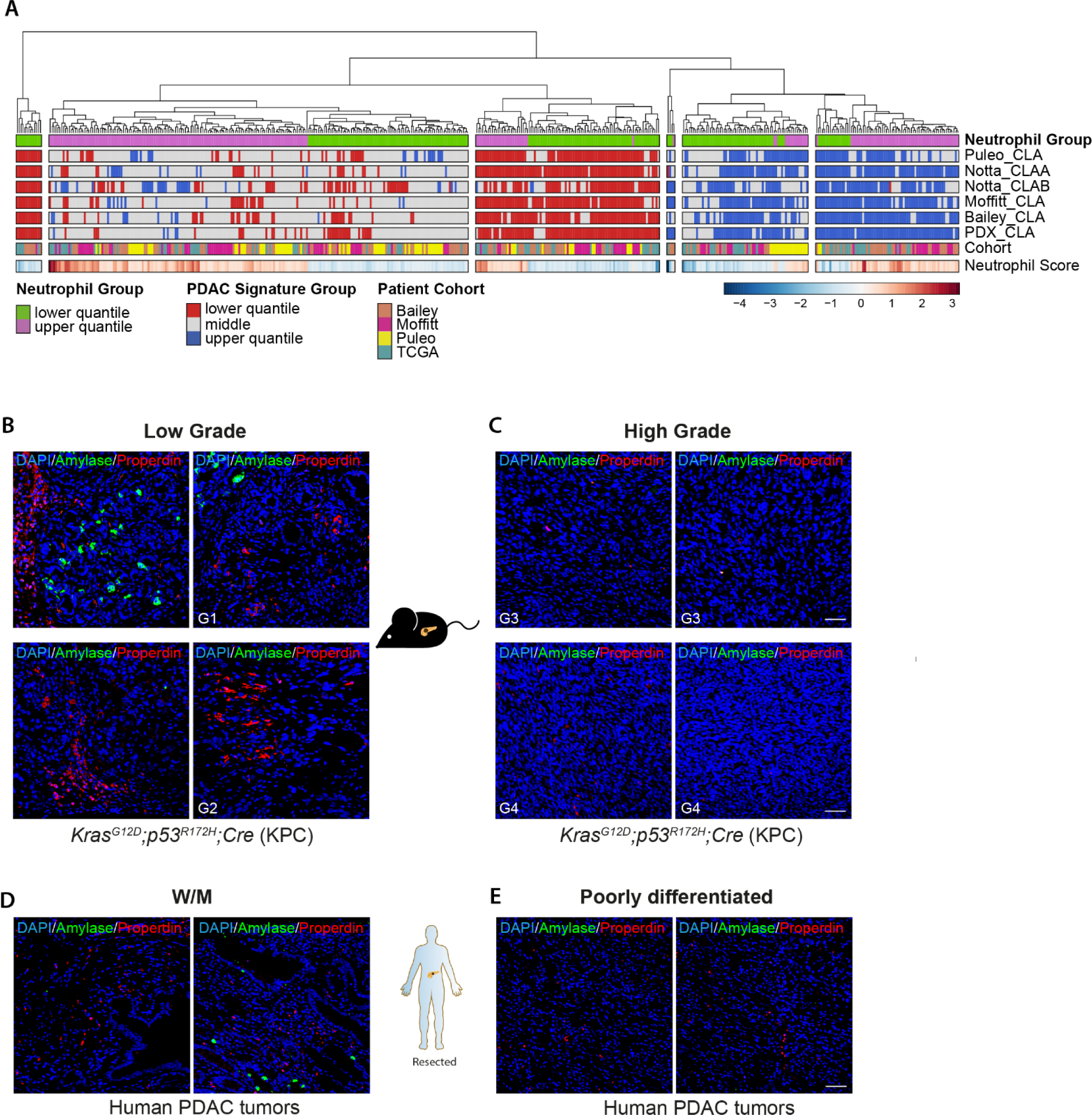
Subtype-specific expression of properdin in PDAC. A) Publicly available data of the Bailey, Moffitt, Puleo and TCGA cohorts with their MCPcounter neutrophil scores in the upper and lower quantile overall (n=234 each) were plotted for neutrophil scores and the indicated CLA-related gene signature Gscores. Annotation of the signatures was analogously based on quantile ranges, higher indicating a more CLA gene expression. For neutrophil scores, row-normalised values are shown. **B** and **C**) Immunofluorescence (IF) staining of amylase (green) and properdin (red) in G1-G2/ low grade (B) and G3-G4/high grade (C) Kras^G12D^;p53^R172H^;Cre (KPC) tumour tissues. **D** and **E**) Properdin expression in primary human PDAC of well-to-moderately (W/M)/CLA (D) and poorly differentiated/BL tumours (E). Scale bars, 50μm. n=5 in low grade/CLA; n=5 in high grade/BL. n=4 in W/M/CLA; n=5 in poorly differentiated/BL tumours.

### Human properdin binds to BL PDAC cells and induces apoptosis

Immunofluorescence microscopy revealed that human properdin bound BL PDAC cell lines, PANC1 (**Fig. 4A**) and MiaPaCa2 (**Fig. 4D**). BL PANC1 was found to bind properdin on the membrane (**Fig. 4A**). Properdin bound BL MiaPaCa2 cells, revealing a “cluster”-like binding pattern on the cell membrane (**Fig. 4A and B**). No FITC was detected in the untreated controls, probed with primary and secondary antibodies, ruling out non-specific properdin binding. Moreover, treatment of the PDAC cell lines with 20 µg/ml of properdin reduced the cell viability of both BL PDAC cell lines by approximately 50% and 70% after 24h and 48h (**Fig. 4C**). A 60% reduction in the cell viability of MiaPaCa2 cells was observed after 48h properdin treatment compared to untreated cells (**Fig. 4D**). No significant difference in cell viability of MiaPaCa2 was observed after 24h treatment (**Fig. 4D**). In addition, apoptosis induction was quantitatively assessed by staining the cells with Annexin V conjugated with FITC + Propidium Iodide (PI) and performing flow cytometry (**Fig. 4E and F**). Nearly 52% reduction was observed in viable PANC1 cells (Cells stained negative for FITC and PI) following treatment with properdin (20 µg/ml) for 24 h (**Fig. 4E**). Similarly, ∼72% reduction of viable PANC1 cells was observed after 48 h treatment ((**Fig. 4E**). In the case of MiaPaCa2 cells, a 50% reduction in viable cells was observed after 48h of properdin treatment and no significant effect was observed in cells treated for 24 h (**Fig. 4F**). Apoptosis induction in PANC1 and MiaPaCa2 following properdin treatment was also confirmed via immunofluorescence microscopy (**Fig. 5A and B**). Fluorescence microscopy of PANC1 (**Fig. 5A**) and MiaPaCa2 (**Fig. 5B**) cells treated with properdin, or the positive control, revealed positive staining for Annexin V (conjugated to FITC), PI and disoriented cell membrane morphology. In contrast, no positive staining of Annexin V or PI was detected in the untreated control.

**Figure 4:**
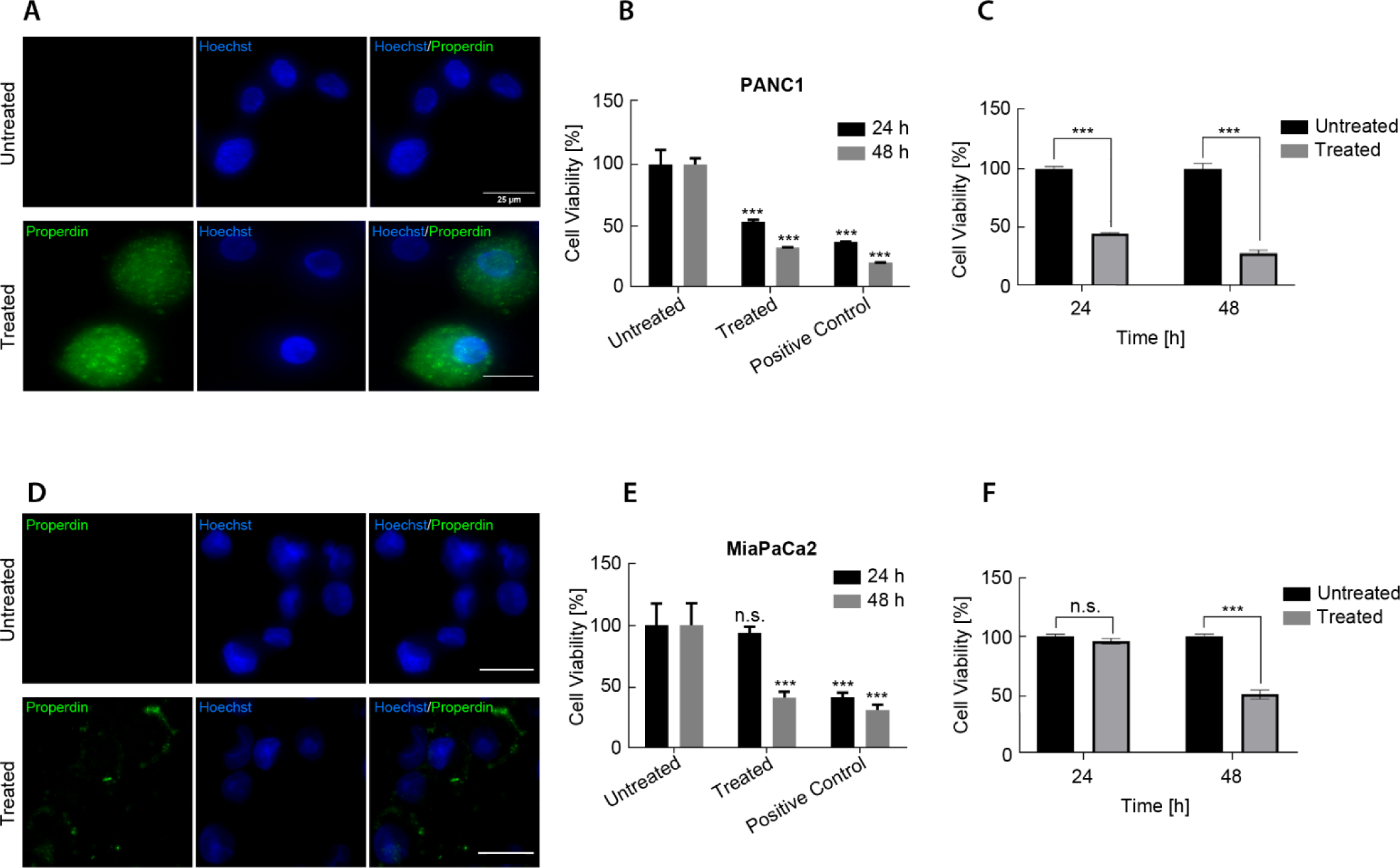
**Properdin binds and induces apoptosis in PDAC cell lines. A and D**) Binding of properdin (20 μg/ml) to BL PDAC cell lines PANC1 (**A)** MiaPaCa2 (D) was studied using fluorescence microscopy. The cell lines were incubated with properdin for 1h at 37 °C followed by 1 h incubation at 4°C. The nucleus of the cells was stained with Hoechst (1:10,000); cells were probed with polyclonal anti-human properdin (1:200) followed by Alexa 488 tagged anti-rabbit IgG. Membrane localisation of the bound proteins was only detected in the treated cells. In contrast, no Alexa 488 signal was detected in the untreated control (cells only). **B** and **E**) MTT assay was performed to assess cell viability. 10,000 PANC1 and MiaPaCa2 cells were treated with properdin (20 µg/ml) for either 24h or 48h. PANC1 **(B)** cell viability reduced by approximately 50% after 24h and ∼70% after 48h of properdin treatment. A 60% reduction in cell viability of MiaPaCa2 **(E)** was observed after 48h properdin treatment compared to untreated cells. No significant difference in cell viability of MiaPaCa2 was observed after 24h treatment. The data is presented as the normalised mean of three independent experiments carried out in triplicates ±SEM. Properdin treated and staurosporine treated positive controls were normalised using the untreated controls (cells + vehicle) as the baseline. Significance was assessed using the two-way ANOVA test (\*\*\**p* < 0.0001). Flow cytometry analysis was performed to determine apoptosis induction in PANC-1 **(C)** and MiaPaCa-2 **(F)** cells treated with 20 μg/ml of properdin. Apoptosis was studied by staining cells with Annexin V/FITC and DNA/PI staining. 12,000 cells were acquired and plotted. The properdin treated cells were normalised using the untreated controls (cells + vehicle) as the baseline. The data is presented as the normalised mean of two independent experiments done in ±SEM. Significance was assessed using the two-way ANOVA test (\*\*\**p* < 0.001).

**Figure 5.**
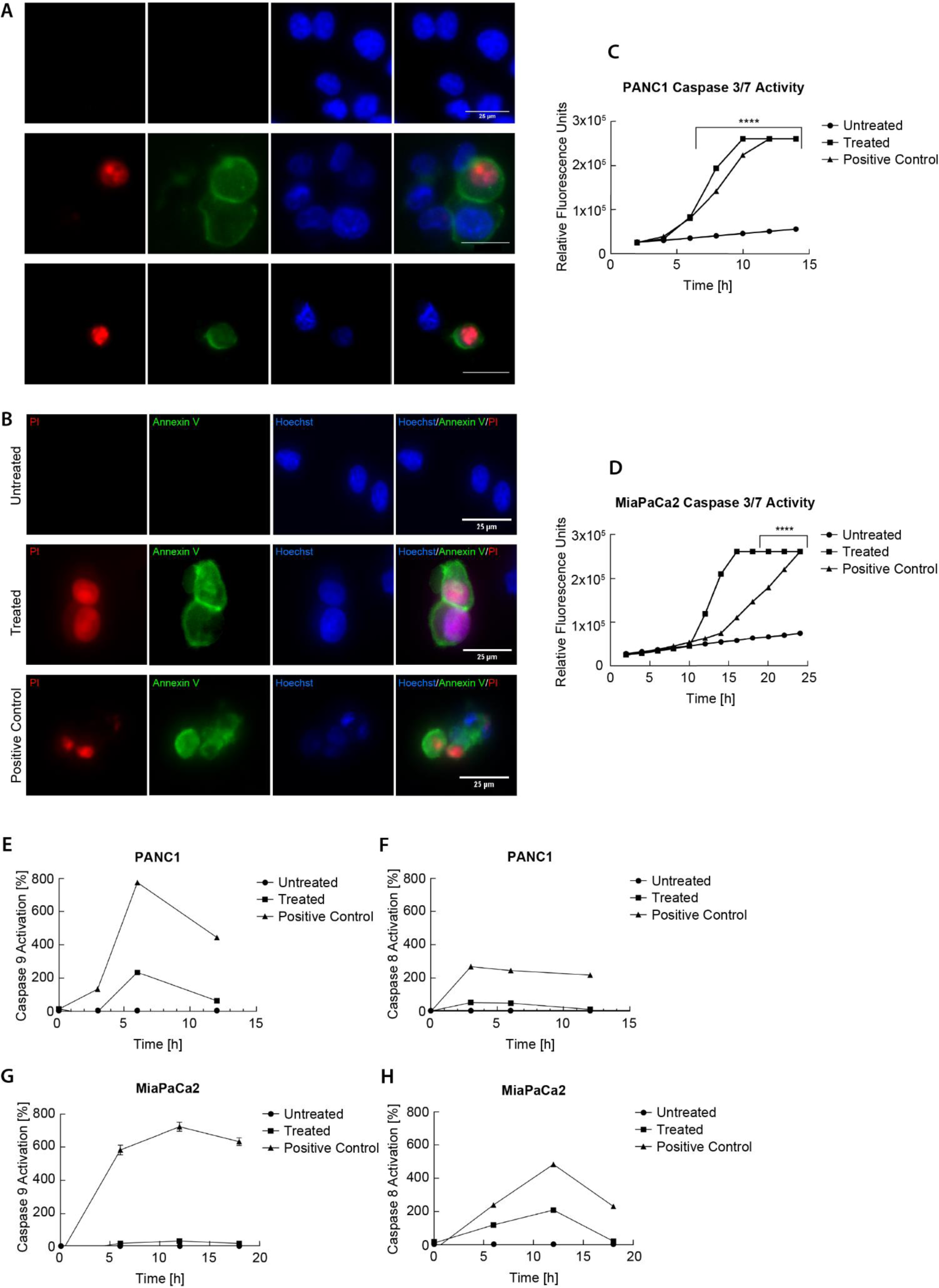
**Properdin treatment induces apoptosis via caspase pathways in PDAC cells**. **A**) Apoptosis induction was analysed using an Annexin V with propidium iodide (PI) staining kit. PANC1 cells **(A)** and MiaPaCa2 cells **(B)** were treated with properdin (20 μg/ml) for 24h and 48h, respectively. The nucleus was stained with Hoechst (1:10,000). The cell membrane of the properdin treated cells and the staurosporine treated positive controls was stained positively with Annexin V/FITC (1:200). The DNA of the treated cells and positive controls were also stained with PI (1:200). No Annexin V/PI staining was detected in untreated cells. **C** and **D**) PANC1 (A) and MiaPaCa2 (B) cells (10,000 cells/well) were treated with properdin (20µg/ml) for 24 h and 48h respectively. The CellEventTM caspase-3⁄7 green detection reagent was used to measure Caspase-3/7 activation. Fluorescence correlating with the activation of caspase-3/7 was recorded using the FITC/Alexa Fluor™ 488 filter settings in the Clariostar plate reader every hour over 48h. Data represent the mean values of three independent experiments. The caspase 3/7 activity in the properdin treated samples, and staurosporine treated positive control was significantly increased compared to the untreated cells (Cells only). Statistical significance was determined using 2-way ANOVA ***P < 0.001. **E**-**H**) PANC1 and MiaPaCa2 (25,000 cells/well) were treated with properdin (20µg/ml) for 15 h and 20h respectively. Caspase-8 and Caspase-9 activity were assayed by the Caspase-Glo® 9 Assay and Caspase-Glo® 8 Assay for PANC1 **(E, F)** and MiaPaCa2 **(G, H)**. Luciferase activity that correlated with the activation of caspase-9 or caspase 8 was recorded using the Clariostar plate reader. The relative caspase activity of properdin treated PDAC cells was calculated using untreated sample (cells + vehicle) as the baseline. Data represent the relative mean value of two independent experiments.

We further examined the likely apoptotic pathway triggered by properdin treatment in PDAC cell lines. Treatment of PANC1 cells with properdin induced caspase 3/7 activity at around 6h time point (**Fig. 5C**). However, properdin treated MiaPaCa2 cells only exhibited caspase 3/7 activity at 10 h (**Fig. 5D**). Furthermore, properdin treatment caused apoptosis in PANC1 cells by inducing caspase 9 activation following 3 to 6h treatment (**Fig. 5E**), while low-level of caspase 9 activity was observed in MiaPaCa2 cells (**Fig. 5G**). In addition, we analysed caspase 8 activity in properdin-treated BL PDAC cells. In the case of MiaPaCa2 cells, properdin treatment triggered the activation of caspase 8, 6h to 12h following treatment with properdin (**Fig. 5H**); however, PANC1 cells treated with properdin also exhibited low-level Caspase 8 activity (**Fig. 5F**). Together, these results suggested that properdin treatment induces cytotoxicity of PDAC cells mainly via caspase 3/7 activation.

## DISCUSSION

Stromal immune components within neoplastic CLA and BL tumours seem to determine prognosis and therapy responsiveness in PDAC patients. The immune system is a critical component of the TME and plays a crucial role in tumour initiation, progression, and metastasis (Ren et al., 2018). Targeting specific immune molecules has been shown to suppress tumour progression and improve chemotherapy effectiveness (Amedei et al., 2014). The most prevalent immune cells within the PDAC TME are macrophages and neutrophils (Nielsen et al., 2021). Tumour-associated macrophages (TAMs) are a crucial inflammatory element of the TME associated with poor prognosis and an unfavourable therapy response in PDAC (Hosein et al., 2019). The abundance of inflammatory cytokines, including IL-1, IL-6, TGF-β, and TNF-α, resulting from high levels of TAMs infiltration, is linked to TME and PDAC progression (Das et al., 2020). A recent study has shown that CCL2, mainly released from BL epithelial cells, induces TAMs to produce TNF-α, leading to highly aggressive PDAC (Tu et al., 2021). Neutrophils are hematopoietic progenitor cells produced in the bone marrow and recruited to the tumour site by cytokines and chemokines produced by the cancer cells (Coffelt et al., 2016). Neutrophils also play a significant role in PDAC and make up a substantial proportion of the immune and inflammatory cells that infiltrate tumours (Jiang et al., 2022a). High levels of neutrophil infiltration in the liver, lung, and stomach occur at the PanIN stage (Jiang et al., 2022a). Neutrophil-derived serum leukotriene B4 (LTB4) has been approved for the early identification of PDAC (Yin et al., 2022). Infiltrating Tumour-Associated Neutrophils (TANs) have unique phenotypes and are divided into N1 and N2 categories: pro-tumorigenic N2 neutrophils and anti-tumorigenic N1 neutrophils (Tyagi et al., 2021). MPO^+^CD11b^+^CD206^-^ and MPO^+^CD11b^+^CD206^+^ are the characteristic cell markers of tumour-associated N1 and N2 neutrophils, respectively (Pu et al., 2019; Tyagi et al., 2021). Studies suggest that N1 neutrophils may have potent anticancer effects through antibody-dependent or direct cytotoxicity, as well as ROS-mediated coupling (Hicks et al., 2006). Infiltrating neutrophils are known to locally secrete properdin, which is stored in their granules (Varghese et al., 2021). Properdin may recognize cancer cells and play a protective role during tumorigenesis, as suggested by some studies (Fuster and Esko, 2005; Sjöblom et al., 2006).

In PDAC, the complement system is a critical factor; surface expression of complement regulatory proteins/receptors, CD46, CD55, and CD59, are well established in PDAC cell lines (Ravindranath and Shuler, 2007). This may lead to complement evasion and limit the effectiveness of novel immunotherapeutic approaches using monoclonal antibodies, which often rely on complement-mediated cytotoxicity (Juhl et al., 1997). Additionally, TGF-β, which is associated with PDAC progression and tissue desmoplasia, inhibits C3 secretion in PDAC cell lines (Andoh et al., 2000). In advanced PDAC patients receiving gemcitabine and intravenous omega-3 fish oil, the restoration of mannose-binding lectin-mediated complement activity via the lectin pathway is linked to a better prognosis (Arshad et al., 2014). Furthermore, the impact of native complement proteins in various types of cancers has been studied. C1q has been described to have anti-tumor properties by inducing apoptosis in prostate, breast, and neuroblastoma, through the activation of WWOX, a tumor suppressor gene, which sends signals that inhibit proliferation and promote apoptosis (Hong et al., 2009; Bandini et al., 2016). In ovarian cancer, C1q uses its globular domains to induce apoptosis through TNF-α and Fas (Kaur et al., 2016). However, in melanoma, C1q promotes tumor cell proliferation, migration, and metastasis (Bulla et al., 2016). In addition, C1q present in the TME of asbestos-induced malignant pleural mesothelioma (MPM), when in conjunction with hyaluronic acid, has been shown to enhance adhesion, migration and proliferation of malignant cells (Agostinis et al., 2017). Factor H has mostly been studied in lung and Cutaneous squamous cell carcinoma, where it protects tumor cells from complement-mediated cytotoxicity and promotes cell migration (Riihilä et al., 2014). However, in mice deficient in Factor H, spontaneous hepatic tumour develops, suggesting its role in controlling unwanted complement activation and avoiding complement-mediated chronic inflammation in the liver (Laskowski et al., 2020). In breast cancer cells, properdin has been found to suppress tumor growth by inducing endoplasmic reticulum stress, independent of complement activation (Block et al., 2019).

In this study, we examined the importance of properdin in PDAC. The differences in the mRNA and protein levels of properdin between normal tissue and PDAC using the UALCAN platform via TCGA and the CPTAC dataset was initially evaluated. A significant increase in properdin gene expression in PDAC was detected compared to the normal pancreas. Furthermore, significant properdin protein expression levels were observed in primary tumours compared to normal tissues. Hence, the tumour purity was examined to assess if the properdin detected in PDAC was secreted/produced by the PDAC cells or by another component of the TME. TIMER v2.0 analysis was performed using the TCGA Pancreas dataset. TIMER is a comprehensive resource for a systematic analysis of immune infiltrates across diverse cancer types. The TIMER v2.0 analysis revealed that CFP expression exhibited a significantly negative correlation with tumour purity, whereas it showed a significant positive correlation with infiltrating levels of neutrophils.

According to PDAC microarray data, most PDAC patients with elevated properdin expression and having increased neutrophil infiltration trended toward higher overall and disease-free survival. Patients with either high neutrophil scores but low CFP expression, or patients with low neutrophil scores and low CFP expression, exhibited a trend toward a lower overall and disease-free survival. The microarray analysis also revealed that patients with a higher neutrophil infiltration were more likely to correlate with the CLA subtype identity. This was further confirmed by immunofluorescent staining of KPC mice and human PDAC tissues. The staining of the PDAC tissues indicated that properdin was primarily generated in low grade or W/M tumours versus high grade or poorly differentiated PDAC tissues. Additionally, Kaplan–Meier survival analysis revealed that properdin gene expression positively correlated with the overall survival rate of CLA and BL type PDAC patients. Furthermore, the cumulative survival analysis performed for CFP expression and levels of infiltrating neutrophils in PDAC revealed that higher levels of infiltrating neutrophils with low CFP expression correlated with a poorer prognosis. However, the analysis also revealed that there is no statistically significant difference in the overall survival in patients with high CFP expression irrespective of neutrophil infiltration levels. Hence, the effect of exogenous properdin against the aggressive BL PDAC cell lines, PANC-1 and MiaPaCa-2, was studied *in-vitro*. The treatment was found to induce apoptosis in the tested PDAC cell lines. We have shown that properdin can bind, PANC-1 and MiaPaCa-2, and reduce cell viability by inducing apoptosis in both the cell lines tested via distinct apoptotic pathways. The activation of the inducer caspase 9 was followed by the start of the executioner caspase 3/7 activation of the in the BL PDAC cells post-treatment. The binding ligand(s) for properdin on the surface of PDAC cells remains to be ascertained.

This study evaluated the effects of properdin on PDAC’s using UALCAN, survival analysis, microarray analysis and apoptosis assays. We report here that increased neutrophil infiltration and local properdin production is associated with improved overall and disease-free survival. Furthermore, properdin was largely found in low grade tumours or W/M/CLA PDAC subtype. In addition, we show that properdin trigger apoptosis in PDAC cell lines PANC-1 and MiaPaCa-2 via distinct pathways. Although these in vitro observations help establish the anti-tumorigenic role of properdin against PDAC cell lines, it is of importance to replicate these possibilities in humanised murine models to mimic the TME of the PDAC. These *in vivo* studies would help elucidate any unknown interaction of properdin with TME components such as hyaluronic acid that could interfere with the anti-tumorigenic activity of properdin. Additionally, it would also be crucial to identify the triggers involved in properdin-induced apoptosis in the PDAC cell lines to help develop properdin as a treatment against PDACs. Furthermore, it would be interesting to examine the specific thrombospondin repeats (TSR) of properdin that may be involved in the anti-tumour activity observed against PDACs.

This study puts neutrophils, together with its ability to secrete properdin, in the PDAC TME at the heart of a range of burning questions in the field. It is clear that innate immunity plays key roles in PDAC since T cells are mostly excluded from its TME. Together with macrophages, neutrophils seem major players here. How the balance between tumour-suppressive N1 and tumour-promoting N2 is maintained and what affects their transition, together with their abundance, could be a consideration for patient stratification in precision immunotherapy. Since TNF-α plays such an important role in subtype switching and CLA/BL heterogeneity, together with their coexistence, the ability of TME neutrophils to produce TNF and properdin differentially can be an important issue in progression versus suppression of PDAC.

## ACKNOWLEDGEMENT

This study was funded by the UAEU (Grant Number-12F043 to UK), and Deutsche Krebshilfe (70112999; 70115054; Max-Eder Program), Fritz-Thyssen Stiftung (1842610), and Wilhelm-Sander-Stiftung (2021.159.1) to SKS. We thank to Frederike Penz for providing technical support.

---

**Supplementary Figure 1.**
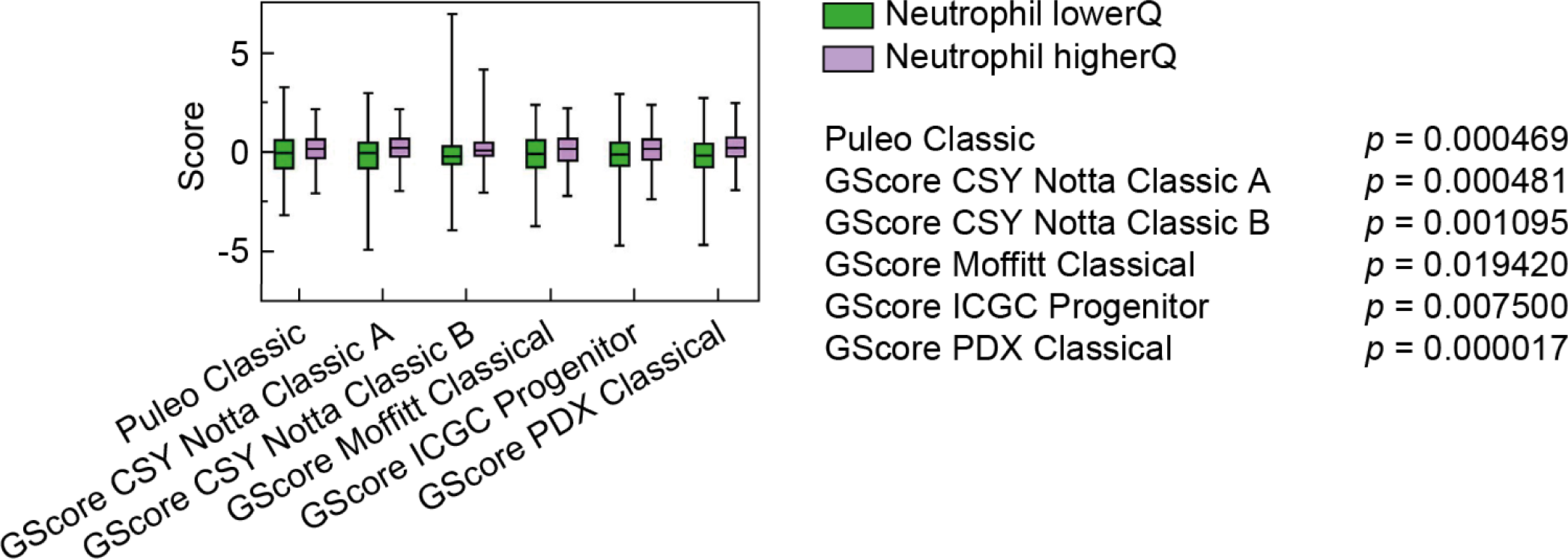
Enrichment scores in PDAC patient sequencing data (TCGA, Moffitt, Bailey, Puleo citation) for indicated subtype signatures in upper and lower quartile of patients based on MCPcounter neutrophil score. Box (25th to 75th percentile with median) and whiskers (min to max) are shown. Unpaired Student’s t-test. Lower quartile, n=234; upper quartile, n=234.

